# IFITM3 deficient mice as a model for testing influenza virus vaccines

**DOI:** 10.1101/2025.05.26.656177

**Authors:** Adrian C. Eddy, Samuel Speaks, Emily A. Hemann, Jacob S. Yount

## Abstract

Influenza virus infections remain a significant global health concern. Development of a universal influenza vaccine has been met with challenges, in part due to difficulties with preclinical vaccine testing in mice, which are widely available but are often poorly infected with human and avian influenza viruses. Here, we investigate whether mice lacking interferon-induced transmembrane protein 3 (IFITM3), an antiviral restriction factor, provide a suitable preclinical model for vaccine testing since we observe enhanced replication of multiple influenza virus strains in these mice. We find that IFITM3 KO mice produce a blunted antibody response to intramuscular vaccination that is increased by a booster dose. Nonetheless, their adaptive immune responses to previous infections and vaccinations were found to be functional in limiting morbidity or viral replication in challenge studies. Overall, our findings identify IFITM3 KO mice as an accessible, functionally immunocompetent preclinical model for assessment of influenza vaccines.

## Introduction

Influenza virus infections cause over 200,000 hospitalizations and more than 10 billion dollars in direct medical costs annually in the US ^1^. In addition to seasonal influenza, the emergence of pathogenic pandemic strains from animals remains a constant concern ^2, 3^. Recent outbreaks of H5N1 avian influenza virus in dairy cattle and poultry have disrupted these industries and sporadic human cases have been reported ^4^. Vaccination remains our best defense against influenza virus, yet seasonal vaccines have gaps in population-level uptake, vary in effectiveness from year to year, and provide minimal protection against emergent strains ^5, 6^. Additionally, virus mutations that circumvent vaccine-mediated immunity highlight the need for novel vaccines which would maintain efficacy despite seasonal mutations and potentially target a wider range of viral subtypes ^7^.

A major limitation in influenza vaccine research is the lack of tractable small animal models for testing new vaccine formulations. Although mice are the most widely used model in preclinical studies, many standard laboratory strains are poorly susceptible to most influenza viruses unless they have been mouse-adapted ^8^. Moreover, mouse adaptation could alter immunogenic sites, including within the HA protein, which is a primary target for antibodies ^9, 10^. Likewise, mouse models with increased susceptibility to human-isolated influenza viruses (*e*.*g*., IFNAR KO, SCID), often lack major components of the immune system, rendering them suboptimal for studying adaptive immune responses ^11-13^.

We have extensively studied interferon-induced transmembrane protein 3 (IFITM3) and developed IFITM3 KO mice on a pure C57BL/6 background ^14-19^ to model deleterious single nucleotide polymorphisms (SNPs) in the human *IFITM3* gene that are associated with increased severity of influenza virus infections ^18, 20-25^. One of these *IFITM3* SNPs was linked to a slightly diminished influenza vaccine response on average, though nearly all the individual antibody responses were within the normal range seen in control subjects ^26^. Moreover, IFITM3 KO mice also showed a delayed peak influenza vaccine antibody responses when vaccinated intraperitoneally ^26^. However, it remains unclear whether IFITM3 KO mice experience altered antibody responses following natural infection or vaccination via the more clinically relevant intramuscular route. Additionally, whether the small defects reported for antibody responses in IFITM3-deficient humans and mice have functional consequences during challenge infections have not been examined.

Here, we examined IFITM3 KO mice for their ability to develop functionally protective adaptive immune responses following infection or intramuscular vaccination with influenza virus HA antigen. We observe that primary vaccine antibody responses are blunted in IFITM3 KO mice as previously reported, but this reduction is largely normalized following booster vaccination. Further, we observe significantly decreased viral titers in vaccinated IFITM3 KO compared to WT upon challenge infection, indicating that a functionally effective adaptive immune response can be generated in the absence of IFITM3. Given the broadly enhanced susceptibility of IFITM3 KOs to influenza viruses, our results suggest that these mice are potentially useful as a pre-clinical influenza vaccine testing model.

## Materials and Methods

### Mice and Vaccinations

IFITM3 KO mice were generated by our lab previously ^17^. WT control animals (Charles River Laboratories, Strain 027) and KO mice were given 2 µg of HA protein (Sino Biological, Cat 40145-V08H1) via 50 µL intramuscular injections. HA protein was combined at a 1:1 ratio with AddaVax adjuvant (InvivoGen, Cat vac-adx-10) or AddaVax and PBS for mock vaccination.

### Viruses and Infections

Influenza viruses used include A/WSN/1933 (H1N1, termed WSN) and the X31 reassortment virus with HA/NA segments from A/Aichi/2/1968 (H3N2) and all remaining segments from A/PR/8/1934 (H1N1) (provided by Dr. Thomas Moran, Icahn School of Medicine at Mount Sinai). A/Victoria/361/2011 (H3N2) and B/Brisbane/33/2008 were acquired from BEI resources (Cat NR-44022 and NR-42006). A/American Green-Winged Teal/Ohio/17OS1834/2017 (H10N8) and A/Black Duck/Tennessee/17OS036/2017 (H5N1) were provided by Dr. Andrew Bowman of the Ohio State University. Viruses were propagated in 10-day embryonated chicken eggs (AVSbio, Cat 10100331) and titered by TCID50 assay on MDCK cells (BEI, Cat NR-2628). Mice were infected intranasally under isoflurane anesthesia. All experiments used female WT and IFITM3 KO mice. Mice losing more than 30% body weight were humanely euthanized. All procedures were approved by the OSU IACUC.

### Antibody Measurements

ELISAs for IgG subtypes were performed as previously described ^27^. The following biotinylated secondary antibodies to IgG subtypes were used: Southern Biotech, Cat 1030-08 (Total IgG), 1070-08 (IgG1), 1090-08 (IgG2B), 1079-08 (IgG2C), 1100-08 (IgG3). Micro-neutralization and hemagglutination inhibition assays of animal serum was performed as previously described ^28^.

## Results

### IFITM3 KO mice mount a protective immune response following low dose IAV infection

To investigate whether IFITM3 KO mice can mount a functional adaptive immune response, we characterized their responses to homologous or heterosubtypic virus rechallenges. We infected IFITM3 KOs with a low dose (200 TCID50) of WSN (H1N1) influenza virus, which caused modest weight loss (**Fig 1A**). Hemagglutination inhibition and neutralizing antibody assays on serum collected at day 18 post infection (p.i.) demonstrated production of influenza virus-specific antibodies in infected IFITM3 KO animals (**Fig 1B-C**). The mice were subsequently challenged with a lethal dose (2,000 TCID50) of WSN IAV on day 30 p.i.. Virus-naive mice rapidly lost weight and died by day 8 p.i., while previously infected mice experienced no appreciable weight loss and demonstrated 100% survival (**Fig 1D-E**), demonstrating that IFITM3 KO mice can develop a robust IAV antibody response and are protected against homologous re-infection.

**Figure 1:**
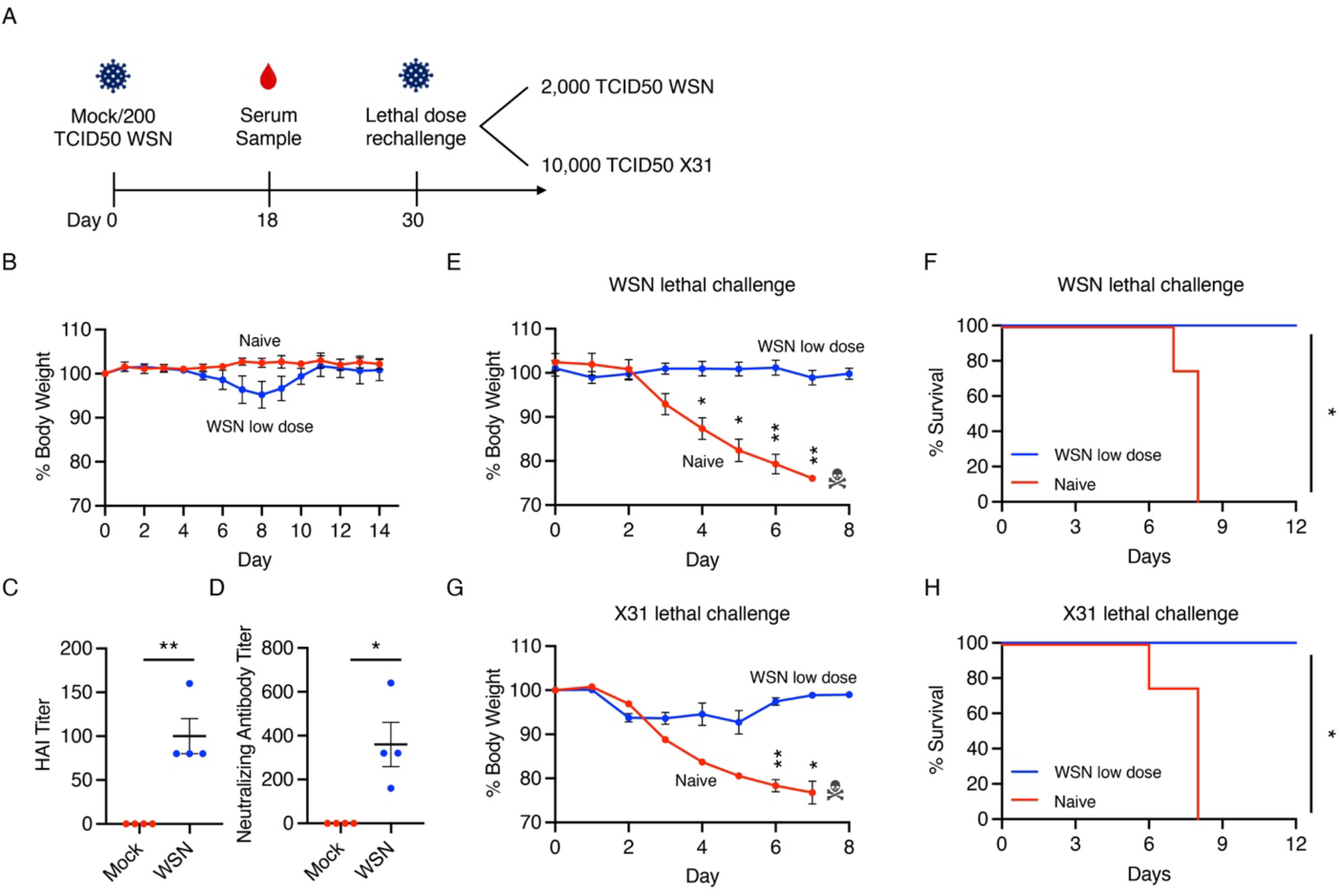
IFITM3 KO mice mount a protective adaptive immune response following live IAV infection. **A-C** IFITM3 KO mice were intranasally infected with 200 TCID50 IAV strain WSN (WSN low dose) or mock infected with saline (Naive). **A** Weight loss measurement for infected and mock-infected animals (each dot is an average of individual mouse weights normalized to 100% relative to day 0, error bars indicate SEM). **B** Hemagglutination inhibition assay with serum from animals in (**A**) using 1% chicken RBC and 100 TCID50 WSN (each dot represents the reciprocal of the highest dilution of animal serum which prevented hemagglutination, error bars indicate SEM, **p<0.01 by unpaired t-test). **C** Micro-neutralization assay with serum from animals in (**A**) using MDCK cells infected and 100 TCID50 WSN (each point represents the reciprocal of the highest dilution of serum which inhibited infection, *p<0.05 by unpaired t-test). **D** On day 30 p.i., animals as in (**A**) were re-infected with a lethal dose of 2000 TCID50 WSN and monitored for weight loss daily (error bars represent SEM, *p<0.05, **p<0.01 by two-way ANOVA followed by Bonferroni multiple comparisons test). **E** Survival analysis of animals in (**D**) (*p<0.05 by Log-rank Mantel-Cox test). **F** On day 30 p.i., another cohort of animals as in (**A**) were re-infected with 10,000 TCID50 X31 and monitored daily for weight loss along with age-matched mock infected mice (error bars represent SEM, *p<0.05, **p<0.01 by two-way ANOVA followed by Bonferroni multiple comparisons test). **G** Survival analysis of animals in (**F**) (*p<0.05 by Log-rank Mantel-Cox test).

We next challenged mice previously infected with the low dose H1N1 WSN with a lethal dose (10,000 TCID50) of the H3N2 reassortant virus commonly referred to as X31 30 days post initial infection. Virus-naive mice rapidly lost weight and succumbed by day 8 p.i. while previously H1N1-infected mice displayed significantly less weight loss in response to the H3N2 X31 virus and achieved 100% survival (**Fig 1F-G**). Taken together, these findings suggest that IFITM3 KO mice have functional immune responses following live virus infection that are protective against subsequent homologous or heterosubtypic virus challenges.

### Non-adapted influenza viruses exhibit enhanced replication in IFITM3 KO mice

Given that most mouse models are minimally susceptible to most influenza viruses, we sought to assess the susceptibility of our IFITM3 KO mouse model to a range of non-adapted influenza viruses. WT and IFITM3 KO mice were infected with 10,000 TCID50 of human influenza viruses A/Victoria/361/2011 (H3N2) or B/Brisbane/33/2008 (Victoria lineage) or with avian influenza viruses A/American Green-Winged Teal/Ohio/17OS1834/2017 (H10N8) or A/Black Duck/Tennessee/17OS036/2017 (H5N1) to measure viral replication in the lungs. IFITM3 KO mice exhibited significantly higher viral replication compared to WT mice for all human and avian strains (**Fig 2A-D)**, demonstrating that IFITM3 KOs allow a greater dynamic range in viral replication for potentially assessing vaccine responses.

**Figure 2:**
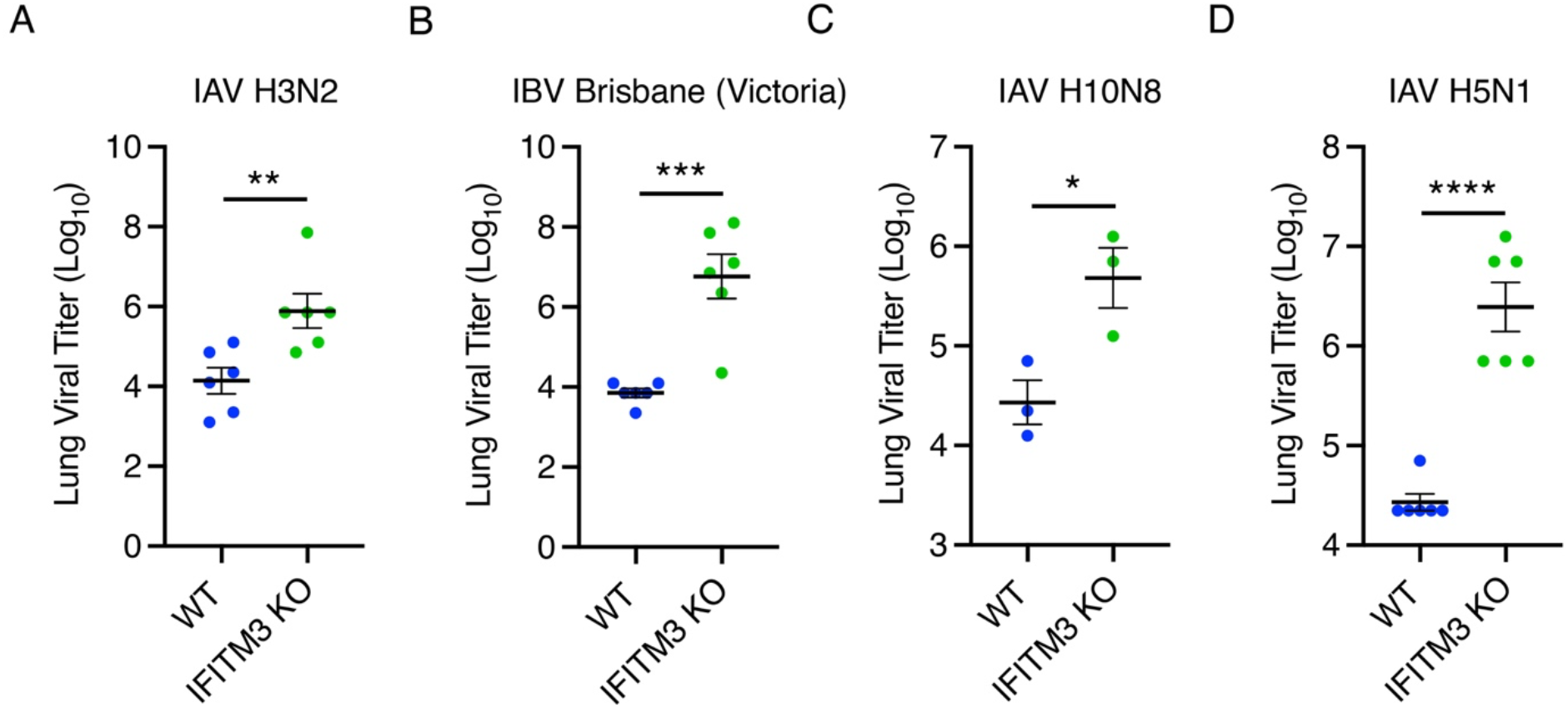
Non-adapted human- and avian-origin influenza viruses exhibit enhanced replication in IFITM3 KO mice. **A-D** Lung viral titer measurements from WT or IFITM3 KO mice sacrificed 5 days p.i. with 10,000 TCID50 of Influenza A/Victoria/361/2011 (H3N2) (**A**), B/Brisbane/33/2008 (Victoria lineage) (**B**), A/American Green-Winged Teal/Ohio/17OS1834/2017 (H10N8) (**C**), or A/Black Duck/Tennessee/17OS036/2017 (H5N1) (**D**). Error bars represent SEM, *p<0.05, **p<0.01, ***p<0.001, ****p<0.0001 by unpaired t-test.

### Vaccination induces HA-specific antibodies in IFITM3 KO mice

WT and IFITM3 KO mice were intramuscularly vaccinated with purified hemagglutinin (HA) from influenza A/Victoria/361/2011 (H3N2) virus with AddaVax adjuvant. Serum was collected following prime and boost vaccinations to evaluate antibody responses (**Fig 3A**). Vaccinated WT mice showed significantly increased levels of HA-specific total IgG, IgG1, and IgG2B antibodies compared to mock-vaccinated controls, while IFITM3 KO mice only achieved significantly increased antibody levels for total IgG and IgG1. Compared to primed WT mice, primed IFITM3 KO mice had significantly lower levels of total IgG, IgG1, and IgG2B, with a non-significant reduction in IgG2C levels, compared to WT controls (**Fig 3B-F**). After the boost, these differences partially normalized, though IgG2 subtypes remained significantly lower in vaccinated IFITM3 KO mice compared to vaccinated WT mice (**Fig 3B-F**). These results confirm that IFITM3 KO mice mount an antibody response to vaccination that is somewhat blunted in comparison to WT mice, though differences are diminished following a boost vaccination.

**Figure 3:**
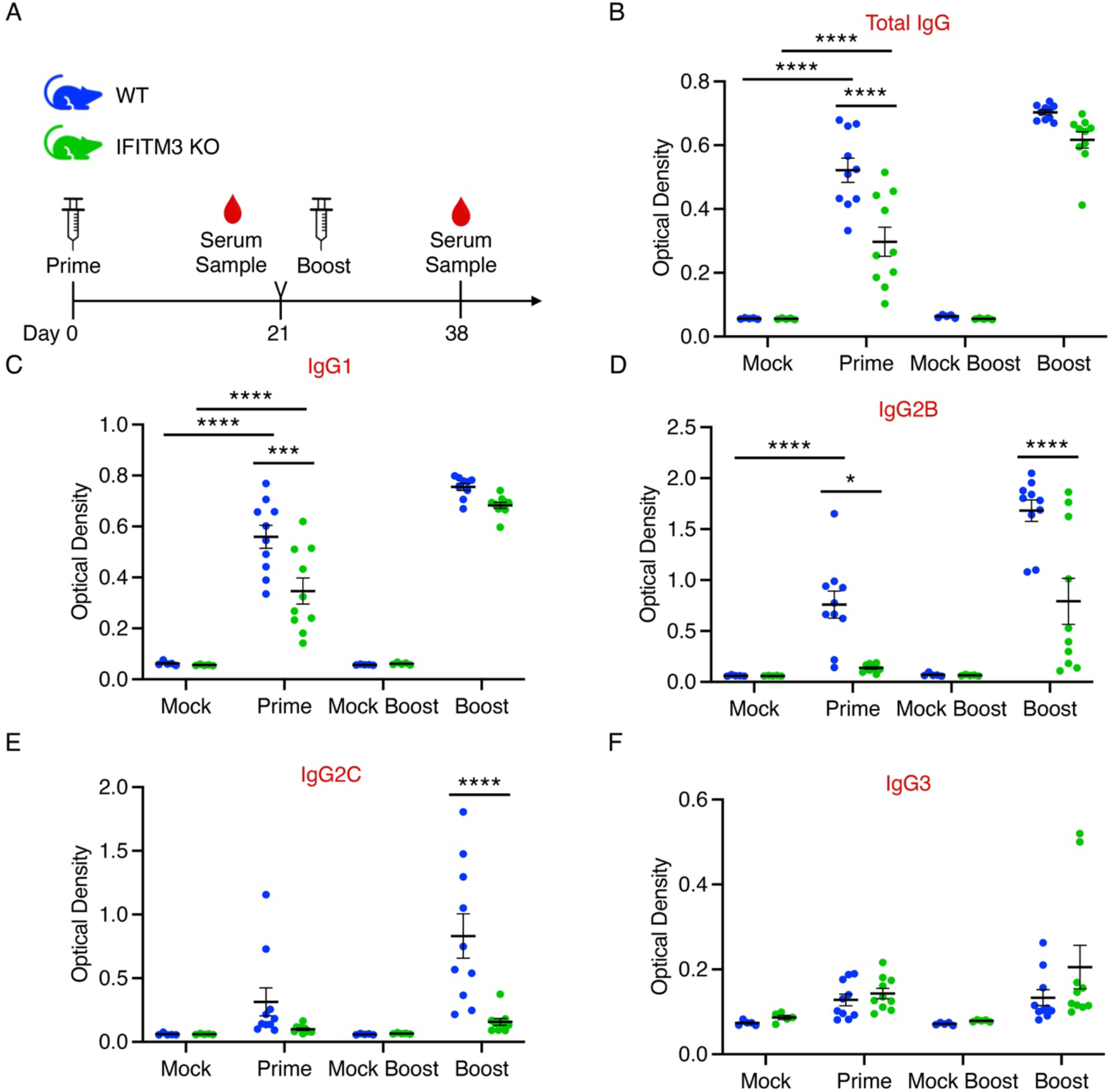
IFITM3 KO mice develop IgG antibodies in response to HA vaccination. **A** Schematic of the HA-based IAV prime-boost vaccination strategy for WT and IFITM3 KO mice (color scheme applies to **B-F**). **B-F** Relative amounts of indicated IgG antibody in WT and IFITM3 KO mouse serum in vaccinated and mock-vaccinated animals measured by ELISA (error bars represent SEM, *p<0.05, **p<0.01, ***p<0.001, ****p<0.0001 by two-way ANOVA followed by Tukey’s multiple comparisons test).

### Vaccinated IFITM3 KO mice develop specific and protective adaptive immune responses

To evaluate the functionality of the antibody response to vaccination in IFITM3 KO mice, neutralizing antibody titers were assessed using serum from prime/boost-vaccinated and mock-vaccinated animals (**Fig 4A**). All vaccinated mice displayed a significantly higher H3N2 neutralizing antibody titer than their mock-treated control counterparts. Within the boost-vaccinated groups, there was no difference in H3N2 neutralization titer between WT and IFITM3 KO mice (**Fig 4B**). Boosted mice were challenged with 10,000 TCID50 of homologous H3N2 IAV, and lung viral titers were measured on day 5 p.i. Both WT and KO vaccinated animals had lower viral loads on average. Notably, only the IFITM3 KO vaccinated group showed a statistically significant reduction compared to mock controls, likely due to the higher baseline viral replication in unvaccinated KO mice, which provided a greater dynamic range for observing vaccine effects (**Fig 4C**). These results indicate that IFITM3 KO mice mount an effective vaccine-induced immune response capable of limiting viral replication.

**Figure 4:**
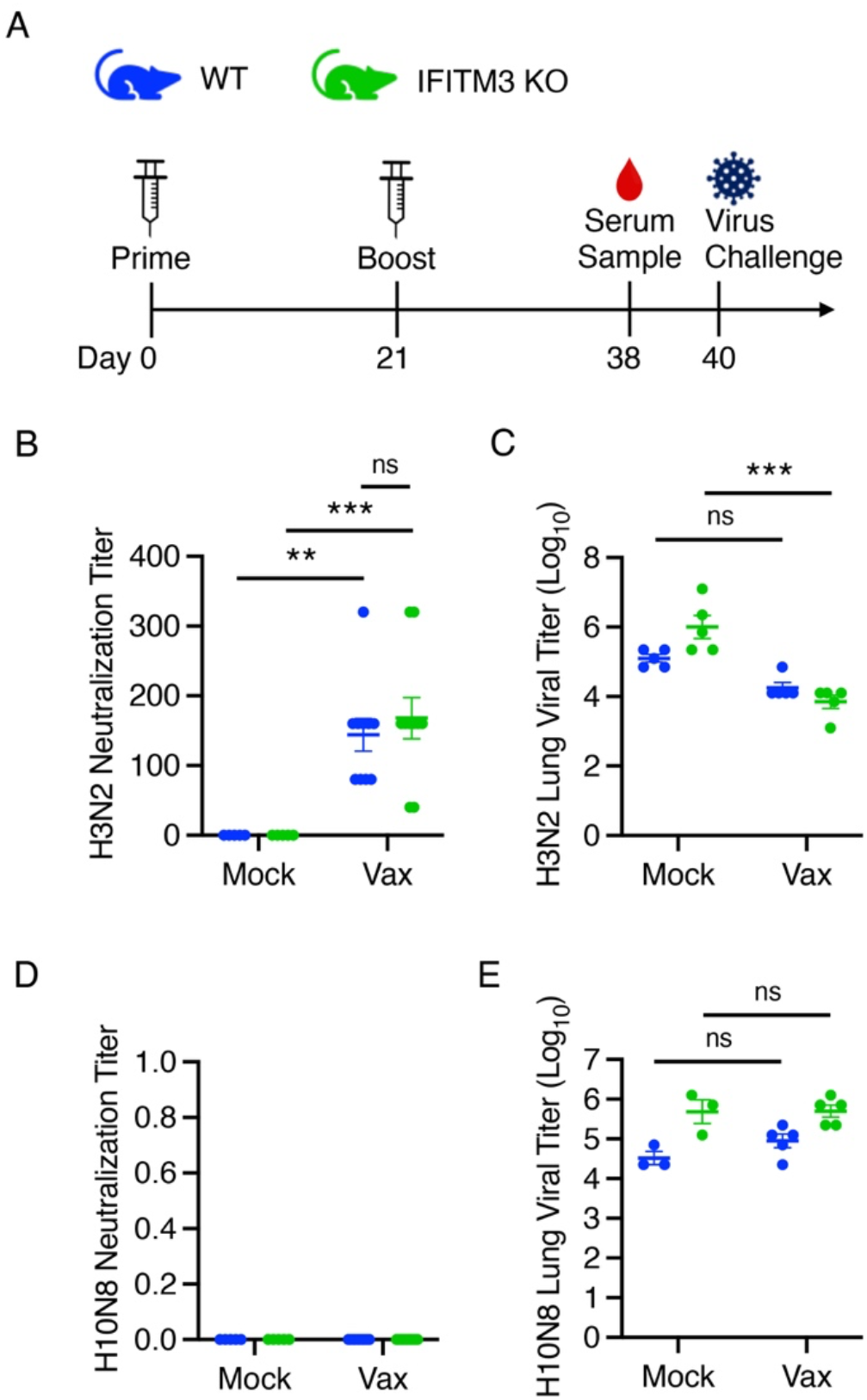
IFITM3 KO mice develop a specific and protective adaptive immune response following HA-based IAV vaccination. **A** Schematic of the HA-based IAV prime-boost vaccination strategy for WT and IFITM3 KO mice (color scheme applies to **B-E**). **B** Micro-neutralization assay was performed with serum from boost-vaccinated and mock-vaccinated mice, added to MDCK cells infected with 100 TCID50 WSN (error bars indicate SEM, **p<0.01, ***p<0.001 by two-way ANOVA followed by Tukey’s multiple comparisons test). **C** Lung viral titers from mice in (**B**) intranasally infected with 10,000 TCID50 H3N2 (error bars represent SEM, ***p<0.001 by two-way ANOVA followed by Tukey’s multiple comparisons test). **D** Micro-neutralization assay was performed as in **A** with 100 TCID50 H10N8. **E** Lung viral titers from mice as in **D** infected with 10,000 TCID50 H10N8 (error bars indicate SEM, p>0.05 by two-way ANOVA followed by Tukey’s multiple comparisons test).

To assess the specificity of this response, we evaluated H10N8 neutralization using serum from HA(H3)-vaccinated mice. No H10N8 inhibition was detected in either WT or KO samples (**Fig 4D**). Additionally, no difference in lung viral titers was observed in vaccinated versus control mice when challenged with H10N8 virus (**Fig 4E**). These findings collectively demonstrate that IFITM3 KO mice can generate a specific and protective adaptive immune response following influenza vaccination.

## Discussion

Influenza virus poses a significant global health burden ^29, 30^, yet preclinical vaccine testing is hindered by the lack of small animal models that support robust viral replication and adaptive immunity. For instance, WT mice are poorly infected by most non-adapted influenza viruses isolated from humans or avian species. Immunodeficient mouse models may allow for higher virus replication, but they often fail to mount a representative adaptive immune response, making them unsuitable for preclinical vaccine testing ^8, 11-13^.

We previously showed that IFITM3 KO mice exhibit increased susceptibility and severity of influenza virus infection^17, 31, 32^. Here we tested whether this model could be used to evaluate vaccine-induced adaptive immunity. While prior studies on IFITM3-deficient humans and mice have reported inconsistent effects on antibody responses ^26, 33, 34^, we observed that IFITM3 KO mice develop functionally effective adaptive immune responses following both infection and vaccination. Initial antibody responses to intramuscular vaccination were blunted in KO mice, consistent with previous reports^26^, but were largely restored after boosting. In protection studies, both homologous and hetrosubtypic challenge infections were controlled, suggesting that B and T cell memory responses are functionally sufficient in the absence of IFITM3. Importantly, IFITM3 KO mice supported higher replication of non-adapted human and avian influenza viruses compared to WT controls. This broader susceptibility allows for a wider dynamic range in lung viral titers for evaluating vaccine efficacy, particularly for strains representative of those currently circulating in humans or zoonotic strains.

Compared to specific-pathogen-free C57BL/6 mice, which often display exaggerated vaccine responses ^35^, IFITM3 KO mice may provide more moderate and thus a more stringent and relevant model for vaccine testing. While “dirty” mouse models have also been suggested to more accurately replicate human immune phenotypes, they are difficult to generate and maintain ^36^. In contrast, IFITM3 KO mice are readily available, genetically defined, and display slightly dampened B cell responses, potentially offering a simpler yet informative platform for early vaccine testing.

In summary, IFITM3 KO mice: 1) permit replication of diverse non-adapted influenza viruses, 2) display increased viral burden allowing for more rigorous evaluation of vaccine efficacy, and 3) exhibit adaptive immune responses to infection and vaccination. These attributes make them a promising tool for rigorous preclinical testing of influenza vaccines. Future studies should extend this model to additional viral pathogens and further explore immune cell functions in the absence of IFITM3.

## Funding

This work was supported by NIH awards R35GM150503 (E.A.H) and R01AI130110 (J.S.Y.).

## Notes

### Competing Interest Statement

The authors have declared no competing interest.

## References

1. Molinari, N.A., et al., The annual impact of seasonal influenza in the US: measuring disease burden and costs. Vaccine, 2007. 25(27): p. 5086–96.

2. Fedson, D.S., Influenza, evolution, and the next pandemic. Evol Med Public Health, 2018. 2018(1): p. 260–269.

3. Bailey, E.S., et al., The continual threat of influenza virus infections at the human-animal interface: What is new from a one health perspective? Evol Med Public Health, 2018. 2018(1): p. 192–198.

4. NCIRD. H5 Bird Flu: Current Situation. 2025 January 18, 2025]; Available from: https://www.cdc.gov/bird-flu/situation-summary/index.html.

5. Taubenberger, J.K. and J.C. Kash, Influenza virus evolution, host adaptation, and pandemic formation. Cell Host Microbe, 2010. 7(6): p. 440–51.

6. Jester, B., et al., Historical and clinical aspects of the 1918 H1N1 pandemic in the United States. Virology, 2019. 527: p. 32–37.

7. Estrada, L.D. and S. Schultz-Cherry, Development of a Universal Influenza Vaccine. J Immunol, 2019. 202(2): p. 392–398.

8. Bouvier, N.M. and A.C. Lowen, Animal Models for Influenza Virus Pathogenesis and Transmission. Viruses, 2010. 2(8): p. 1530–1563.

9. Gitelman, A.K., et al., Changes in the antigenic specificity of influenza hemagglutinin in the course of adaptation to mice. Virology, 1984. 134(1): p. 230–2.

10. Hirst, G.K., Studies on the Mechanism of Adaptation of Influenza Virus to Mice. J Exp Med, 1947. 86(5): p. 357–66.

11. Shepardson, K.M., et al., IFNAR2 Is Required for Anti-influenza Immunity and Alters Susceptibility to Post-influenza Bacterial Superinfections. Front Immunol, 2018. 9: p. 2589.

12. Wu, H., et al., Sustained viral load and late death in Rag2-/-mice after influenza A virus infection. Virol J, 2010. 7: p. 172.

13. McNab, F., et al., Type I interferons in infectious disease. Nat Rev Immunol, 2015. 15(2): p. 87–103.

14. Yount, J.S., et al., Palmitoylome profiling reveals S-palmitoylation-dependent antiviral activity of IFITM3. Nat Chem Biol, 2010. 6(8): p. 610–4.

15. Yount, J.S., R.A. Karssemeijer, and H.C. Hang, S-palmitoylation and ubiquitination differentially regulate interferon-induced transmembrane protein 3 (IFITM3)-mediated resistance to influenza virus. J Biol Chem, 2012. 287(23): p. 19631–41.

16. Chesarino, N.M., T.M. McMichael, and J.S. Yount, Regulation of the trafficking and antiviral activity of IFITM3 by post-translational modifications. Future Microbiol, 2014. 9(10): p. 1151–63.

17. Kenney, A.D., et al., IFITM3 protects the heart during influenza virus infection. Proc Natl Acad Sci U S A, 2019. 116(37): p. 18607–18612.

18. Zani, A. and J.S. Yount, Antiviral Protection by IFITM3 In Vivo. Curr Clin Microbiol Rep, 2018. 5(4): p. 229–237.

19. Chesarino, N.M., T.M. McMichael, and J.S. Yount, E3 Ubiquitin Ligase NEDD4 Promotes Influenza Virus Infection by Decreasing Levels of the Antiviral Protein IFITM3. PLoS Pathog, 2015. 11(8): p. e1005095.

20. Everitt, A.R., et al., IFITM3 restricts the morbidity and mortality associated with influenza. Nature, 2012. 484(7395): p. 519–23.

21. Pan, Y., et al., IFITM3 Rs12252-C Variant Increases Potential Risk for Severe Influenza Virus Infection in Chinese Population. Front Cell Infect Microbiol, 2017. 7: p. 294.

22. Wang, Z., et al., Early hypercytokinemia is associated with interferon-induced transmembrane protein-3 dysfunction and predictive of fatal H7N9 infection. Proc Natl Acad Sci U S A, 2014. 111(2): p. 769–74.

23. Allen, E.K., et al., SNP-mediated disruption of CTCF binding at the IFITM3 promoter is associated with risk of severe influenza in humans. Nat Med, 2017. 23(8): p. 975–983.

24. Kenney, A.D., et al., Human Genetic Determinants of Viral Diseases. Annu Rev Genet, 2017. 51: p. 241–263.

25. Zhang, Y.H., et al., Interferon-induced transmembrane protein-3 genetic variant rs12252-C is associated with severe influenza in Chinese individuals. Nat Commun, 2013. 4: p. 1418.

26. Lei, N., et al., IFITM3 affects the level of antibody response after influenza vaccination. Emerg Microbes Infect, 2020. 9(1): p. 976–987.

27. Hemann, E.A., et al., A Small Molecule RIG-I Agonist Serves as an Adjuvant to Induce Broad Multifaceted Influenza Virus Vaccine Immunity. J Immunol, 2023. 210(9): p. 1247–1256.

28. Cuevas, F., et al., An In Vitro Microneutralization Assay for Influenza Virus Serology. Curr Protoc, 2022. 2(7): p. e465.

29. Preventing Seasonal Flu. 2024 [cited 2025 January 19]; Available from: https://www.cdc.gov/flu/prevention/index.html.

30. Buchy, P. and S. Badur, Who and when to vaccinate against influenza. Int J Infect Dis, 2020. 93: p. 375–387.

31. Kenney, A.D., et al., Influenza virus replication in cardiomyocytes drives heart dysfunction and fibrosis. Sci Adv, 2022. 8(19): p. eabm5371.

32. Denz, P.J., et al., Innate immune control of influenza virus interspecies adaptation via IFITM3. Nat Commun, 2024. 15(1): p. 9375.

33. Qin, L., et al., High Level Antibody Response to Pandemic Influenza H1N1/09 Virus Is Associated With Interferon-Induced Transmembrane Protein-3 rs12252-CC in Young Adults. Front Cell Infect Microbiol, 2018. 8: p. 134.

34. Lee, J., et al., IFITM3 functions as a PIP3 scaffold to amplify PI3K signalling in B cells. Nature, 2020. 588(7838): p. 491–497.

35. Coughlan, L., Caught in a trap: How pre-clinical studies in laboratory mice exaggerate vaccine responses. Cell Rep Med, 2021. 2(12): p. 100484.

36. Fiege, J.K., et al., Mice with diverse microbial exposure histories as a model for preclinical vaccine testing. Cell Host Microbe, 2021. 29(12): p. 1815–1827 e6.

